# Prenatal Exposure to Environmentally Relevant Low Dosage Dibutyl Phthalate Reduces Placental Efficiency in CD-1 Mice

**DOI:** 10.1101/2024.02.26.582170

**Authors:** Tasha Pontifex, Xinran Yang, Ayna Tracy, Kimberlie Burns, Zelieann Craig, Chi Zhou

## Abstract

**Introduction:** Dibutyl phthalate (DBP), a phthalate congener, is widely utilized in consumer products and medication coatings. Women of reproductive age have a significant burden of DBP exposure through consumer products, occupational exposure, and medication. Prenatal DBP exposure is associated with adverse pregnancy/fetal outcomes and cardiovascular diseases in the offspring. However, the role of fetal sex and the general mechanisms underlying DBP exposure-associated adverse pregnancy outcomes are unclear. We **hypothesize** that prenatal DBP exposure at an environmentally relevant low dosage adversely affects fetal-placental development and function during pregnancy in a fetal sex-specific manner.

**Methods:** Adult female CD-1 mice (8-10wks) were orally treated with vehicle (control) or with environmentally relevant low DBP dosages at 0.1 μg/kg/day (refer as DBP0.1) daily from 30 days before pregnancy through gestational day (GD) 18.5. Dam body mass composition was measured non-invasively using the echo-magnetic resonance imaging system. Lipid disposition in fetal labyrinth and maternal decidual area of placentas was examined using Oil Red O staining.

**Results:** DBP0.1 exposure did not significantly affect the body weight and adiposity of non-pregnant adult female mice nor the maternal weight gain pattern and adiposity during pregnancy in adult female mice. DBP0.1 exposure does not affect fetal weight but significantly increased the placental weight at GD18.5 (indicative of decreased placental efficiency) in a fetal sex-specific manner. We further observed that DBP0.1 significantly decreased lipid disposition in fetal labyrinth of female, but not male placentas, while it did not affect lipid disposition in maternal decidual.

**Conclusions:** Prenatal exposure to environmentally relevant low-dosage DBP adversely impacts the fetal-placental efficiency and lipid disposition in a fetal sex-specific manner.

## Introduction

Phthalates are a group of environmental endocrine disruptors that widely exist in the environment and adversely impact cardiovascular and metabolic function in humans^1-5^. Although general usage of phthalates has gradually reduced in the past 5-10 years, dibutyl phthalate (DBP), a phthalate congener, is still widely utilized in consumer products and medication coatings^3, 6-9^. Despite decreasing trends in legacy phthalate use in response to enhanced awareness and regulation, a strong need for phthalate research remains as reductions in phthalate burden differ by age, socio-economic, and geographical groups for specific phthalate metabolites^10-13^. Women of reproductive age have a significant burden of DBP exposure^3, 6, 7^ through consumer products, occupational exposure, and medication (0.1-233µg/kg/day)^8, 9^. Higher urinary DBP metabolite levels (indicative of higher exposure) were observed in Hispanic and non-Hispanic black women than in other ethnic groups^14^, strongly suggesting disparities among women of different backgrounds.

Multiple biomonitoring studies reported DBP exposure in most mothers and babies during pregnancy^15-17^. Prenatal DBP exposure is associated with adverse pregnancy/fetal outcomes (e.g., fetal growth restriction)^15-17^, childhood obesity^17, 18^, and cardiovascular diseases in the offspring^5, 19-22^. Studies in rats have shown that prenatal DBP exposure at 500mg/kg/day leads to reduced placental weight^23, 24^. In humans, maternal DBP exposure in 1^st^ trimester is reported to be associated with increased risks of fetal intrauterine growth restriction (which is associated with placental dysfunction) in pregnancies with male but not with female fetuses^25^. However, mechanisms underlying these adverse effects of prenatal DBP exposure are unclear.

Fetal sex has significant impacts on the effects of prenatal stress exposure-associated adverse perinatal and pregnancy outcomes^26-29^. Pregnancies with male fetuses are reported to be at a higher risk of preterm birth, while female fetuses are at a higher risk of fetal growth restriction^27^. Although prenatal exposure to other phthalate congeners (DMP and DEHP) is eported to be associated with fetal sex-specific fetal growth restriction^25^ in humans, the role of fetal sex in DBP exposure-associated adverse pregnancy/fetal outcomes is still unclear.

To date, most DBP studies in the literature evaluated dosages that are much higher than human snvironmental exposure^5, 30^. Furthermore, the majority of the limited existing literature investigating environmentally relevant dosages of DBP exposure focused on the gonad toxicity in non-pregnant mice. Although oral ingestion of environmentally relevant DBP exposure results in detection of its active metabolite in the ovary and disruption of various ovarian processes and pathways (e.g., folliculogenesis, steroidogenesis, xenobiotic metabolism, DNA repair and apoptosis), the effects of environmentally relevant low dosages DBP exposure on pregnancy and fetal development are still not clear.

In this study, we tested the hypothesis that environmentally relevant low dosage DBP exposure differentially dysregulates male and female fetal placental function during pregnancy using an established DBP exposure mouse model^8, 31^. We examined the effect of environmentally relevant DBP exposure on the weight gain and body composition of adult female mice pre-pregnancy and during pregnancy. We further determined the effect of prenatal environmentally relevant low dosage DBP exposure on fetal/placental development as well as placental lipid disposition at gestational day (GD) 18.5.

## Methods

### Ethical Approval

All animal experiments followed guidance provided in the Guide for the Care and Use of Laboratory Animals by National Research Council^32^ and the Animal Research: Reporting of In Vivo Experiments (ARRIVE) guidelines^33^ with the approval of the University of Arizona Institutional Animal Care and Use Committee (Protocol# 20-670).

### Animals

Adult (8-10wks) female CD-1 mice were obtained from Charles River Laboratories (Charles River, California) and housed in single-use, BPA-free cages at the University of Arizona Animal Facility. Water and food were provided ad libitum, their light cycle maintained at 14 L:10 D, and temperature at 22 ± 1°C. After an acclimation period of at least 24 h, mice were randomly assigned to receive oral tocopherol-stripped corn oil (MP Biomedicals, OH; vehicle; control group) or an environmentally relevant^3, 6-9^ DBP (Sigma-Aldrich, MO) dosage (100, 10, and 0.1 μg/kg/day; refer as DBP100, DBP10, and DBP0.1).

### Oral DBP Dosing in Non-pregnant and Pregnant Mice

Determining Environmentally Relevant DBP Dosages That Does Not Affect Weight Gain in Non-Pregnant Adult Female Mice

To determine an environmentally relevant DBP dosages that does not affect the weight gain in non-pregnant female CD-1 mice, adult female CD-1 mice (8-10wks) were orally treated with vehicle (control group; n=14) or with environmentally relevant DBP dosages (DBP100, DBP10, or DBP0.1; n=8 per group) daily for 30 days. Mice were subjected to daily weighing, vaginal smear collections, and oral dosing with vehicle or DBP treatments. Weight of each mouse before the first day of treatment was used as baseline weight to calculate the percentage of weight gained during treatment.

Environmentally Relevant Low-Dosage DBP Dosing in Adult Female Mice Pre-Pregnant through Pregnancy

Based on our dose finding experiment in non-pregnant mice, DBP0.1 did not affect the weight gain in adult female mice. Hence, DBP0.1 was used in subsequent experiments to determine physiological changes that are independent of weight gain. Adult female CD-1 mice (8-10wks) were orally treated with vehicle (control [CT] group; n=18) or DBP0.1 (n=18) daily for 30 days before mating with male CD-1 mice without DBP exposure and throughout pregnancy until gestational day (GD) 18.5. The weight of each mouse was measured daily. Weight of each mouse before the first day of treatment was used as baseline weight to calculate the percentage of weight-gain during treatment. At GD18.5, mice were euthanized under pre-sedation with inhaled isoflurane followed by decapitation. Major maternal organs (heart, lung, liver, kidney, ovary) were collected and weighted at GD18.5. In addition, all fetus and placentas of each dam were collected and weighted at GD18.5.

### Body Mass Composition Measurement

Control and DBP0.1-treated dams were subjected to body mass composition (fat and lean mass) measurement non-invasively while conscious using an echo-magnetic resonance imaging system (EchoMRI-900 body composition analyzer with A10 insert for mice)^34, 35^ at day 0, 15, 30 of pre-pregnancy treatment as well as GD5.5, GD11.5, and GD18.5 during pregnancy.

### Histology and Ovarian Corpora Lutea (CL) Counts

After the end of the experiment, ovaries from each dam were dissected followed by fat and oviductal tissue removal. Both ovaries were fixed for subsequent histological processing. Fixation was done in neutral buffered formalin overnight at 4°C with shaking. After fixing, ovaries were washed in 70% ethanol, processed, and embedded in paraffin. Embedded ovaries were serially sectioned (5μm thickness) throughout the tissue. Two sequential sections were kept and mounted every 100µm to generate various histological evaluation levels throughout the ovary. The resulting sections were processed for hematoxylin (Richard-Allan, 7211) and eosin (Richard-Allan, 7111) staining using a standard procedure. Stained ovary slides were imaged prior to analysis as described^8^. Briefly, each ovarian section was imaged at 10x magnification using a Leica DMIL LED microscope. Ovarian sections too large to fit in one view, were imaged and captured in tiles, and then merged using the stitching tool in Adobe Photoshop. Only fully developed, functional CLs were counted and compared between treatments.

### Oil Red O Staining of Mouse Placenta

Lipid disposition in placenta is closely linked to placental function and efficiency^36, 37^. The neutral lipids in mice placenta tissues were stained using an Oil Red O Staining Kit (IHC World Cat# IW-3008) following the manufacturer’s instruction. In brief, mice placenta tissues were embedded in optimal cutting temperature (OCT) compound and stored in -80°C until sectioning. Frozen mice placenta sections (10μm thick, 2 per individual placenta) were fixed in 10% formalin for 5 mins. After fixing, the placental sections were incubated in Pre-Stain Solution for 5 mins at room temperature followed with 10 min Oil Red O solution incubation at 60°C. After rinsing, nuclei of sections were counterstained with Mayer’s Hematoxylin for 1 min. After rinsing, sections were mounted using Aqueous Mounting Medium, and imaged at 40x magnification using a brightfield microscope (Leica DM5500B). Lipid droplets area in mice placental tissue sections were quantified using National Institutes of Health (NIH) ImageJ software (http://rsbweb.nih.gov/ij/).

### Statistical Analyses

SigmaPlot software (Systat Software., San Jose, CA) was used for statistical analyses. Data are represented as the medians ± standard deviation (SD). Data analyses were performed using the Student’s t-test, Mann-Whitney Rank Sum test, One-way ANOVA, Kruskal-Wallis test, and two-way ANOVA as appropriate. Differences were considered significant when *P* <0.05. Benjamini and Hochberg False Discovery Rate (FDR)-adjustment^38, 39^ was used for multiple comparison correction as appropriate.

## Results

### Effect of DBP exposure at environmentally relevant dosages on the weight gain of non-pregnant female mice

To determine the DBP exposure dosage that does not affect non-pregnant adult female mice, animals were orally treated with vehicle (CT group), DBP100, DBP10, or DBP0.1 daily for 30 days. Compared with control group, 30-day exposure to DBP100 and DBP10 does-dependently increased the weight of non-pregnant female CD-1 mice (Fig.1A). Specifically, female mice with DBP100 exposure significantly gained weight starting after 18 days of treatment (10.8%) and peaked at 20 days of treatment (15.3%). Mice with DBP10 exposure exhibit a trend of weight gain after 20 days of treatment and reach significant weight gain of 5.6%-6.9% by 23 to 25 days after treatment. Similar to the control (CT) group, 30 days of DBP0.1 exposure does not cause significant weight gain of female mice (Fig.1B).

**Figure 1.**
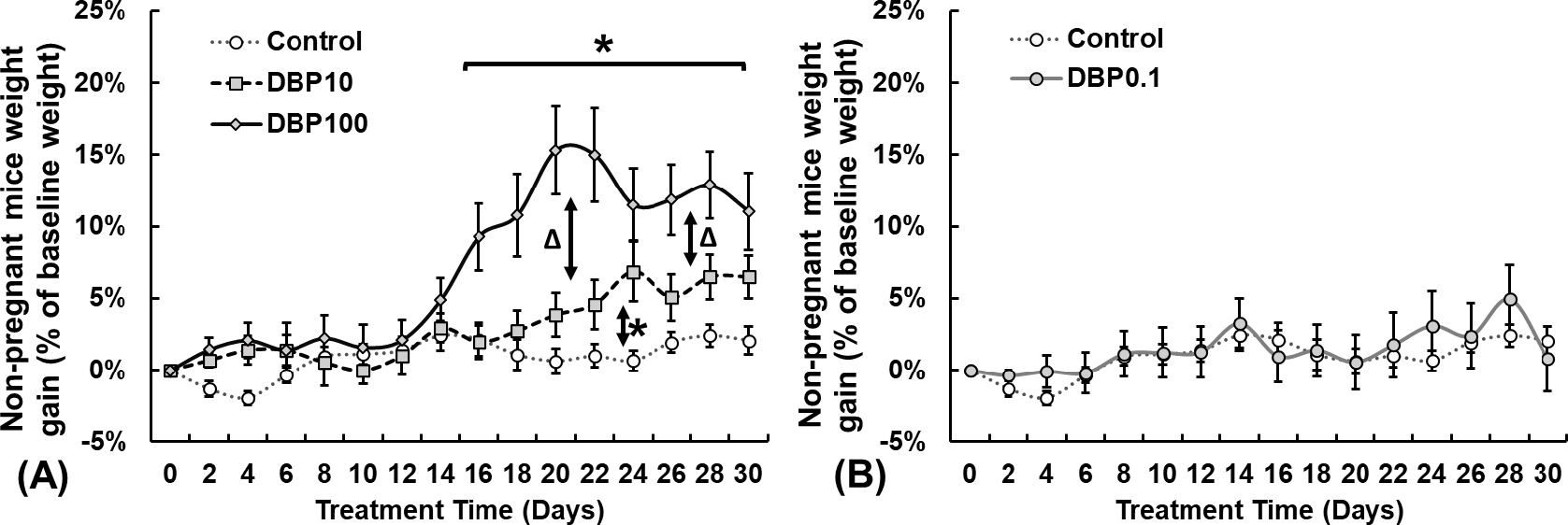
Adult female CD-1 mice weight gain compared to control over 30 days of DBP exposure at (A) 100, 10, and (B) 0.1 μg/kg/day. Data expressed as median ± standard error of mean. DBP100: 100μg/kg/day (n=8); DBP10: 10μg/kg/day (n=8); DBP0.1: 0.1μg/kg/day (n=8); Control: Vehicle only (n=14). **\***: Differ from Control (P<0.05). ^**Δ**^: Differ between DBP100 and DBP10 (P<0.05).

### Effect of DBP0.1 exposure on female fertility efficiency in CD-1 mice

DBP0.1 exposure decreased the pregnancy rate in the female mice from 94.44% to 83.33% (Fig.2A), but it did not affect number of live pups per litter (litter size Fig.2B), male to female fetus ratio (Fig.2C), and ovarian CLs number per dam (Fig.2D) compared to CT group. Interestingly, DBP0.1 exposed mice exhibited significantly lower pup to CL ratio at GD18.5 (defined as number of pups divided by number of CLs, Fig.2E).

**Figure 2.**
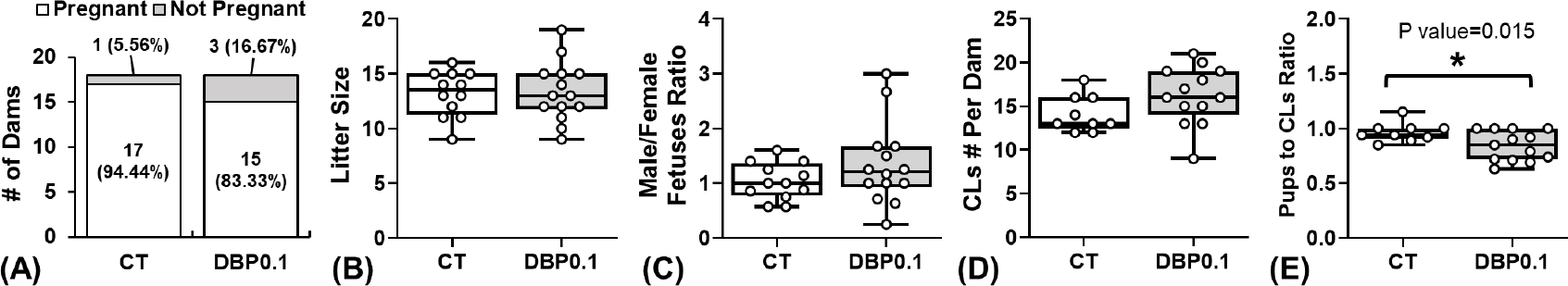
Effect of DBP0.1 exposure on female fertility parameters in control (CT) and DBP0.1 treated CD-1 mice. (A) Pregnancy rate (n=18 for each group), (B) litter size (CT: n=12; DBP0.1: n=14), (C) male/female fetus ratio per litter (CT: n=12; DBP0.1: n=14), and (D) pups to CLs ratio (CT: n=9; DBP0.1: n=13) in CT and DBP0.1 treated CD-1 mice. **\***: Differ between CT and DBP0.1 groups (P<0.05). Each data point represents the data from an individual dam.

### Effect of DBP0.1 exposure on weight gain patterns of pregnant mice

Our daily weight measurement revealed that DBP0.1 exposure does not affect the pre-pregnancy and pregnancy weight gain patterns compared to control group (Fig.3A). Our body composition data (EchoMRI™ LLC, TX; Fig.3B) further showed that DBP0.1 exposure does not significantly alter the body fat percentage of dams (from 30 days pre-pregnancy through GD18.5). Further, DBP0.1 exposure did not affect the weight of the maternal heart, lung, kidney, and liver at GD18.5 compared to control group (Fig.3C-F).

**Figure 3.**
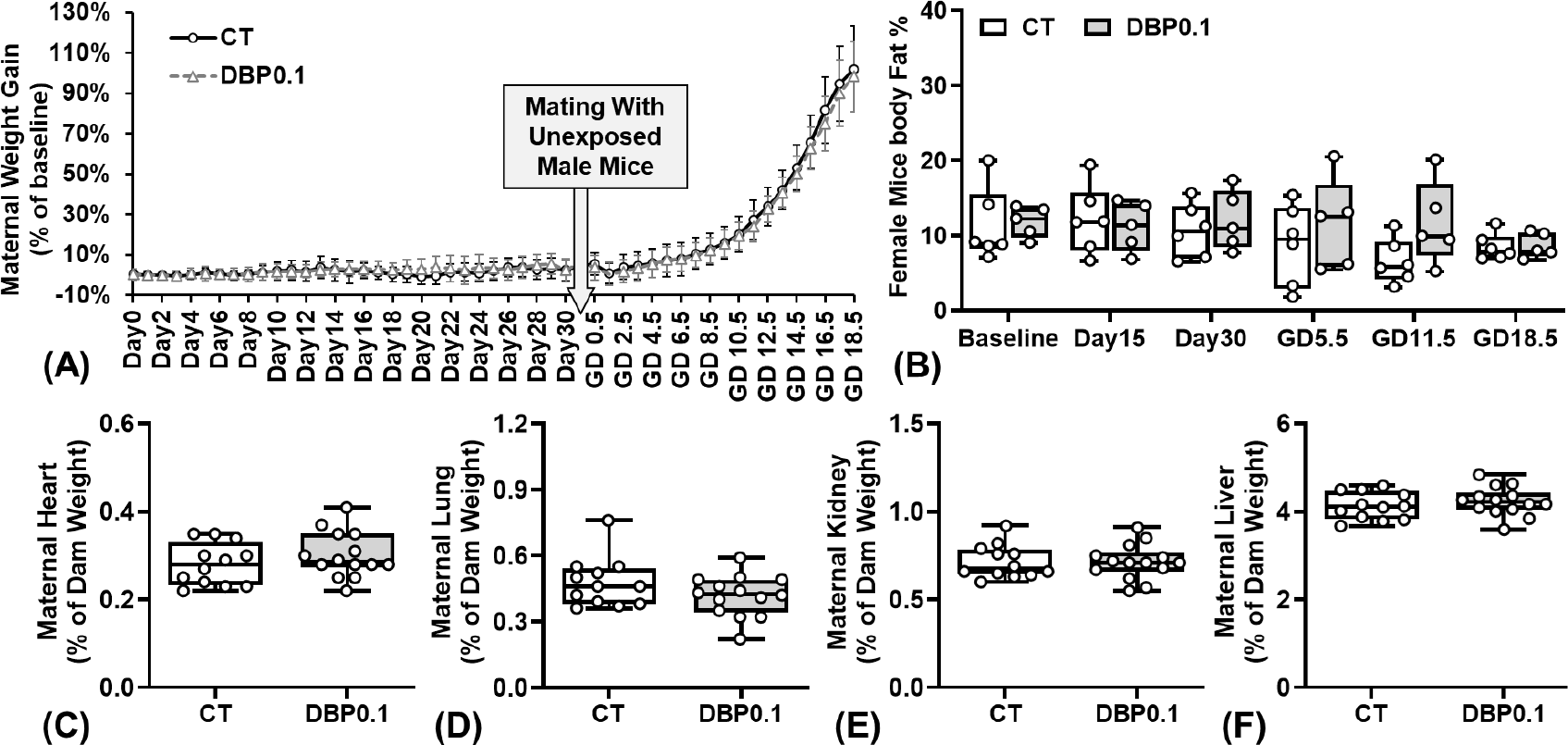
Effect of DBP0.1 exposure on weight gain patterns of pregnant dams. (A) Daily weight gain (CT: n=15; DBP0.1: n=14), (B) adiposity (body fat %; n=6 for each group), and the weight of maternal (C) heart, (D) lung, (E) kidney, (F) liver at GD18.5 in CD-1 mice (C-F: CT group: n=12; DBP0.1 group: n=14). (A) Data expressed as median ± standard deviation. (B-F) Each data point represents the data from an individual dam.

### Effect of Prenatal DBP0.1 exposure on placental efficiency in female and male fetuses

Compared to the CT group, DBP0.1 exposure did not affect fetal weight (Fig.4A) but significantly increased the placental weight (Fig.4B) of both male and female fetuses at GD18.5 (indicative of decreased placental efficiency). There is no difference between the weight of male and female fetuses from both CT and DBP0.1 dams. However, male placental weights are significantly higher than female placentas in CT group. The DBP0.1 exposure-increased placenta weight is more dramatic in female than in male placentas and eliminated the fetal sex-specific placental weight differences observed in CT group.

**Figure 4.**
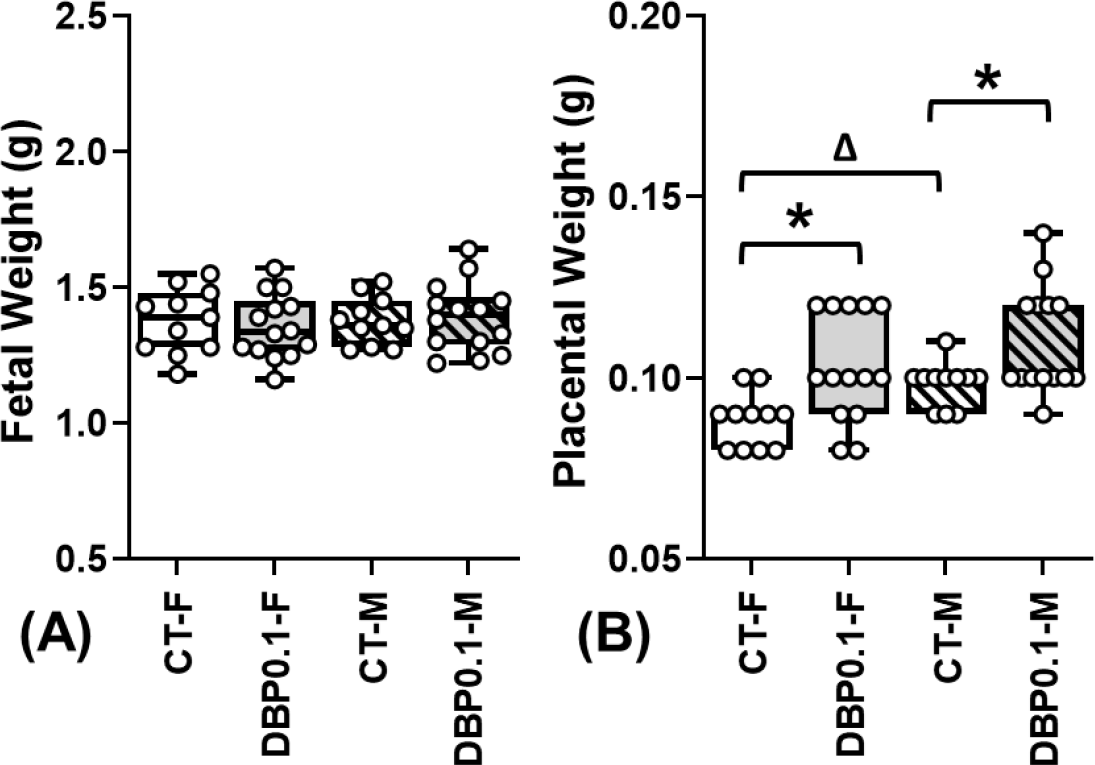
Effect of DBP0.1 exposure on fetal and placental weight at GD18.5. DBP0.1 exposure did not alter fetal weight (A), while it significantly increased placental weight (B) in female (F) and male (M) fetuses in CD-1 mice by GD18.5 (CT-F: n=12; CT-M: n=12; DBP0.1-F: n=14; DBP0.1-M: n=14). *:Differ between CT and DBP0.1 group within the same fetal sex; ^**Δ**^:Differ between F and M fetuses within the same treatment group (P <0.05). Median value of all female or male fetuses from the same dam was shown as a single data point in figures.

### Effect of prenatal DBP0.1 exposure on female and male placental lipid disposition pattern

In placentas from control dams, lipid disposition (calculated as the neutral lipid area relative to tissue area) in the fetal labyrinth area of the placenta is significantly higher in female than in male placentas (Fig.5A). The fetal sex-specific fetal labyrinth lipid disposition pattern observed in control placentas was diminished in placentas from the DBP0.1 group. DBP0.1 exposure significantly reduced the lipid disposition in the fetal labyrinth area of female placentas but did not significantly affect the lipid disposition in male placentas. DBP0.1 exposure does not affect the lipid disposition in the maternal decidual area in both female and male placentas (Fig.5B).

**Figure 5.**
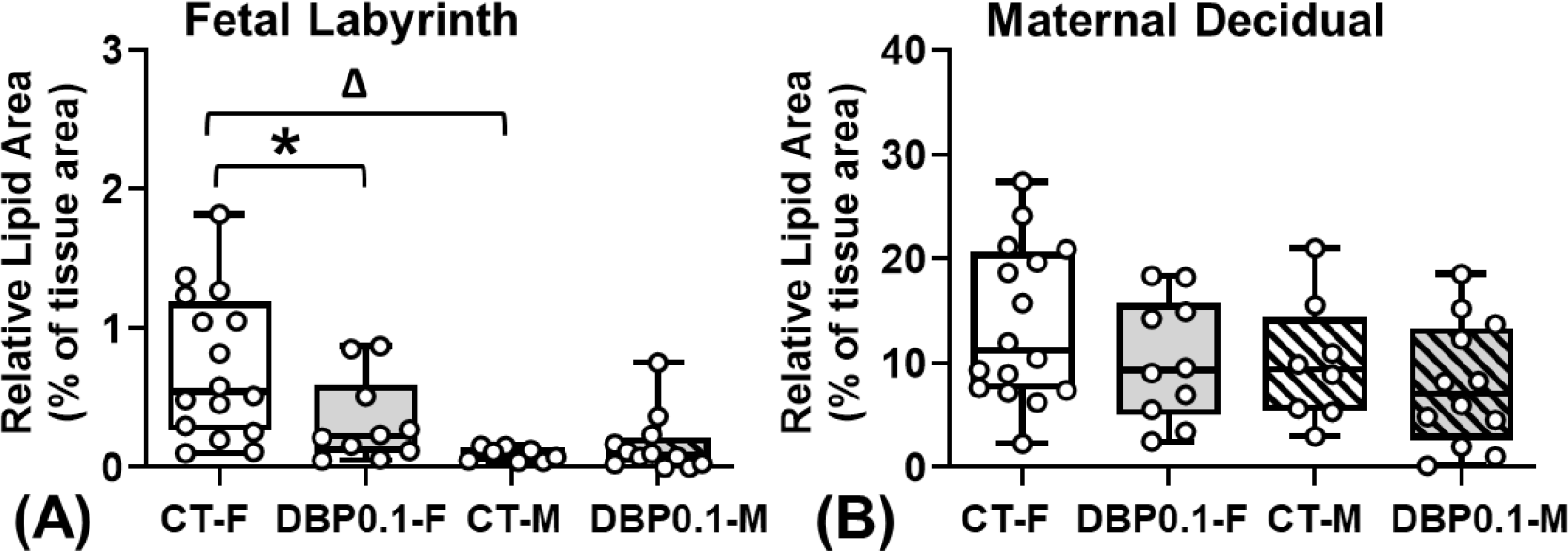
Lipid disposition in (A) fetal labyrinth and (B) maternal decidual of mice placental tissues from CT and DBP0.1 groups. *: Differ between CT and DBP0.1 within the same fetal sex; ^**Δ**^: Differ between female and male fetuses within the same treatment group (P <0.05 Two-way ANOVA). The median value of all images from the same placenta was shown as a single data point (CT-F: n=15; DBP0.1-F: n=10; CT-M: n=8); DBP0.1-M: n=11).

## Discussion

In this study, we have demonstrated that although daily dosing with a low, environmentally relevant exposure to DBP of 0.1 µg/kg/day does not affect the weight (both pre-pregnant and during pregnancies) of adult female CD-1 mice, it significantly increases the placental weight and reduces the placental efficiency at GD18.5. Further, we have shown for the first time that prenatal DBP0.1 (0.1μg/kg/day) exposure caused fetal sex-specific reductions in the fetal labyrinth lipid disposition in the placenta while not affecting the lipid disposition in the maternal decidual area at GD18.5 in CD-1 mice. These data clearly indicate that environmentally relevant low-dosage DBP exposure at DBP0.1 causes fetal sex-specific placental dysfunction.

Consistent with previous reports^8^, we observed that 30 days of environmentally relevant dosages of DBP exposure (DBP100 and DBP10) dose-dependently increased the weight of non-pregnant adult female mice. However, our data showed that 30 days of environmentally relevant DBP exposure at a much lower environmentally relevant dosage at DBP0.1 causes no significant weight gain nor body fat percentage changes in non-pregnant adult female mice compared to control. DBP is known to be associated with obesity^40-42^, metabolic disorders^43^, and cardiovascular diseases^25, 44^ in humans. Maternal obesity is a prevalent risk factor for pregnancy complications (e.g., preterm birth, fetal growth restriction)^45-49^. In studies utilizing higher DBP exposure dosages that cause significant maternal weight gains, it would be very difficult to separate the adverse pregnancy outcomes caused by DBP exposure and by the maternal weight gain caused by DBP exposure. In this study, we specifically chosen an environmentally relevant low DBP dosage (DBP0.1) that does not significantly affect the weight gain in female mice, which allows us to evaluate the effect of maternal exposure to environmentally relevant low-dosage DBP during pregnancy without being affected by the maternal weight gain that would have occurred with higher DBP dosages.

Although the reproductive toxicity of DBP on male fertility is well established^50, 51^, the effect of DBP exposure on female fertility is not fully understood. Existing literature on the effects of DBP exposure on pregnancy outcomes using animal models often utilizes DBP dosages that are much higher than the human environment^5, 30, 52^. Nonetheless, these studies have identified various endpoints of interest including increased offspring adiposity and body weight^52^, and lack of glucose homeostasis^53^. Despite this, the effect of environmentally relevant low-dosage DBP exposure during pregnancy on fetal development and pregnancy outcomes remains elusive.

Our data have shown that although 30 days of DBP0.1 exposure did not affect the body weight or body fat composition of non-pregnant adult female mice, it significantly reduced the reproductive efficiency of the female mice. When mating with untreated CD-1 male mice, the DBP0.1 exposure reduced the pregnancy rate in female CD-1 mice from 94.44% to 83.33%. In addition, although DBP0.1 exposure did not significantly affect the litter size, fetal sex ratio, and ovarian CL count per dam, it significantly reduced the pup-to-CL ratio which is indicative of a disconnect between ovulation and live pups. Although DBP has been shown to disrupt ovarian folliculogenesis in mice^8, 54, 55^, this disconnect between ovulation and pups is likely not due to a defect in ovarian release of eggs but rather due to embryo loss during pregnancy. This possibility is supported by previous reports describing that DBP is an endocrine disruptor that adversely affects pregnancy outcomes^30, 42^. Specifically, human epidemiological studies report associations between phthalate exposures and increased risk of recurrent pregnancy loss^56^; thus, the disconnect between CL number and pups could be due to embryo loss but further studies will be needed to address this possibility directly. Observations in this current study extended the DBP exposure-associated adverse effects on female fertility to an environmentally relevant DBP exposure dosage to as low as 0.1μg/kg/day (DBP0.1).

Our data have shown that DBP0.1 exposure allows for examination of the effect the environmentally relevant DBP exposure during pregnancy independent of the effect of maternal weight gain that would usually cause by higher dosages of DBP. Specifically, adult female mice with 30-days pre-pregnancy DBP0.1 exposure and continued daily DBP0.1 exposure throughout pregnancy did not have significant changes in weight gain pattern and the body fat composition when compared with control animals. Further, this DBP0.1 treatment did not change the weight of major maternal organs at GD18.5. We further observed that the prenatal DBP0.1 exposure did not affect fetal weight but significantly increased placental weight and reduced fetal/placental weight ratio at GD18.5, indicative of reduced placental efficiency in pups from DBP0.1 exposed dams. Further, our data showed that DBP0.1 causes more dramatic placental weight increase in female than male placentas and eliminated the sex dimorphisms of placental weight seen in control animals, which indicates more dramatic reduction in placental efficiency in female fetuses. All these data show that although DBP exposure at an environmentally relevant low dosage (DBP0.1) seems not to affect maternal weight and body composition during pregnancy, it still fetal sex-specifically affects placental development and placental efficiency.

Lipids play a crucial role in placental function^57, 58^. Dysregulation in placental lipid dynamics, as observed in our study, indicates disrupted nutrient transportation and overall placental function^58-61^. We observed a notable sexual dimorphism in fetal labyrinth lipid disposition in placentas from control mice, with female placentas exhibiting significantly higher fetal labyrinth lipid accumulation compared to their male counterparts. Strikingly, DBP0.1 exposure led to a significant reduction in fetal labyrinth lipid disposition in female placentas but does not affect the fetal labyrinth lipid disposition in male placentas. Our observation that DBP0.1 exposure abolishes the fetal sex-specific difference in fetal labyrinth lipid disposition, highlighting the disruptive effects of DBP on normal placental development and function. This is further supported by the fetal sex-specific dysregulation of placental weight in DBP0.1 exposed animals that observed in the current study.

Interestingly, although DBP0.1 exposure leads to fetal sex-specific dysregulation of fetal labyrinth lipid disposition, it does not affect lipid disposition in maternal decidual areas of placental tissues both female and male. This data indicates that the DBP0.1 exposure mainly dysregulates lipid disposition in the fetal labyrinth area of the placentas in a fetal sex-specific manner. This observation aligns with the established role of the labyrinth as the primary nutrient exchange structure in the placenta^62, 63^, where dynamic lipid trafficking and metabolism is essential to sustain fetal growth and development. This data also agrees with previous reports showing that DBP and its metabolite can cross the placenta and play important roles in lipid metabolism and transportation^64-67^.

In conclusion, our data have demonstrated that environmentally relevant low dosage of DBP exposure at 0.1μg/kg/day reduces the reproductive efficiency in female mice and fetal sex-specifically dysregulates the placental efficiency and lipid disposition. Findings in this study highlight the significance of including fetal sex as a biological variable when examining the impact of environmental endocrine disruptor exposures on placental function and fetal development.

### Perspectives

To date, mechanisms underlying adverse effects of prenatal DBP exposure are unclear. Further, most studies in the literature evaluated DBP dosages that are much higher than the human environmental exposure. Here we reported that DBP exposure as low as 0.1μg/kg/day during pregnancy, though does not affect maternal weight gain, still causes fetal sex-specific dysregulation of placental development and fetal labyrinth lipid disposition. These observations highlight the significance of further studying the effect of environmental endocrine disruptor exposure during pregnancy on fetal development at a human environment relevant dosage as well as the necessity to include fetal sex as a biological variable in future investigations.

## Acknowledgments

We would like to thank Dr. Xiaosong Liu for technical assistance.

